# A conserved long-range RNA interaction in SARS-CoV-2 recruits ADAR1 to enhance virus proliferation

**DOI:** 10.1101/2025.07.07.663401

**Authors:** Siwy Ling Yang, Louis DeFalco, Sainan Wang, Yi Hao Wong, Jian Han, Chee Keng Mok, Kiat Yee Tan, Su Ying Lim, Zhiya Zhao, Yu Zhang, Jovi Jian An Lim, Joy S. Xiang, Radoslaw Sobota, Lin-Fa Wang, Justin Jang Hann Chu, Andres Merits, Roland G. Huber, Yue Wan

## Abstract

Long-range RNA-RNA pairing impacts the genome structure and function of SARS-CoV-2 variants. To understand the structure and function relationships of different SARS-CoV-2 variants that have emerged during the COVID-19 pandemic, we performed high-throughput structure probing and modelling of the genomic structures of the wildtype (WT), Alpha, Beta, Delta and Omicron variants of SARS-CoV-2. We observed that genomes of SARS-CoV-2 variants are generally structurally conserved, and that single-nucleotide variations and interactions with RNA binding proteins can impact RNA structures across the viruses. Importantly, using proximity ligation sequencing, we identified many conserved ultra-long-range RNA-RNA interactions, including one that spans more than 17 kb in both the WT virus and the Omicron variant. We showed that mutations that disrupt this 17 kb long-range interaction reduce viral fitness at later stages of its infection cycle, while compensatory mutations partially restore virus fitness. Additionally, we showed that this ultra-long-range RNA-RNA interaction structure binds directly to ADAR1 to alter the RNA editing levels on the viral genome. These studies deepen our understanding of RNA structures in the SARS-CoV-2 genome and their ability to interact with host factors to facilitate virus infectivity.

## INTRODUCTION

SARS-CoV-2 belongs to the family *Coronaviridae* characterized by large (26 – 32 kb) positive-strand RNA genomes^1^. Since its emergence in 2019, SARS-CoV-2 has caused millions of deaths world-wide^2,3^. In addition to the original WT strain, many SARS-CoV-2 variants including Alpha, Beta, Delta and Omicron variants of concern (VOCs) have emerged and circulated around the world^3,4^. Compared to the WT virus, VOCs contain different amounts of mutations along their genome which results in differences in their infectivity among humans. Among the VOCs, the Omicron contains 128 mutations as compared to the WT strain and is the most divergent VOC in terms of its genomic sequence (**Supp. Figure 1a-c**)^5^.

Numerous studies have shown that different RNA structures, including the frameshifting element (FSE), present within the SARS-CoV-2 genome are important for virus functions^6–9^. Several studies devoted to mapping of secondary structure of the SARS-CoV-2 genome, both in infected cells and in virions, have been published since 2020^10–18^. However, the locations of variable and conserved RNA structures across genomes of different VOCs, how single-nucleotide variations (SNVs) and RNA binding proteins (RBPs) impact RNA structures, and how ultra-long-range RNA-RNA pairing within the genome impacts virus functions are still under-explored. Here, we probed the RNA secondary structures of WT, Alpha, Beta, Delta and Omicron variants to identify structurally variable and conserved regions along their genomes. We identified highly conserved regions along the SARS-CoV-2 genome, as well as structural changes due to mutations and RBPs binding in the RNA genome. We also identified conserved ultra-long-range RNA-RNA interactions that span distance more than 10 kb, across the variants, including a paired RNA structure element that spans 17 kb on the SARS-CoV-2 genome. Mutations impairing this interaction and compensatory mutations restoring this long-range interacting RNA structure revealed that it is important for virus fitness. We observed that this element can bind to ADAR1 and induce RNA editing along the virus genome. This study expands our understanding of the structure and function of the SARS-CoV-2 RNA genomes.

## RESULTS

### Mapping of RNA structures across five different variants of SARS-CoV-2

To probe the RNA secondary structures along genomes of SARS-CoV-2 variants, we infected Vero-E6 TMPRSS2 cells using WT, Alpha, Beta, Delta or Omicron variants and treated the cells with a structure probing compound (NAI) at 48 hours post-infection (hpi) (**Figure 1a**). We then extracted the RNAs, performed reverse transcription, library construction and deep sequencing, followed by mapping the reads to the Vero transcriptome as well as to the respective SARS-CoV-2 genomes.

**Figure. 1.**
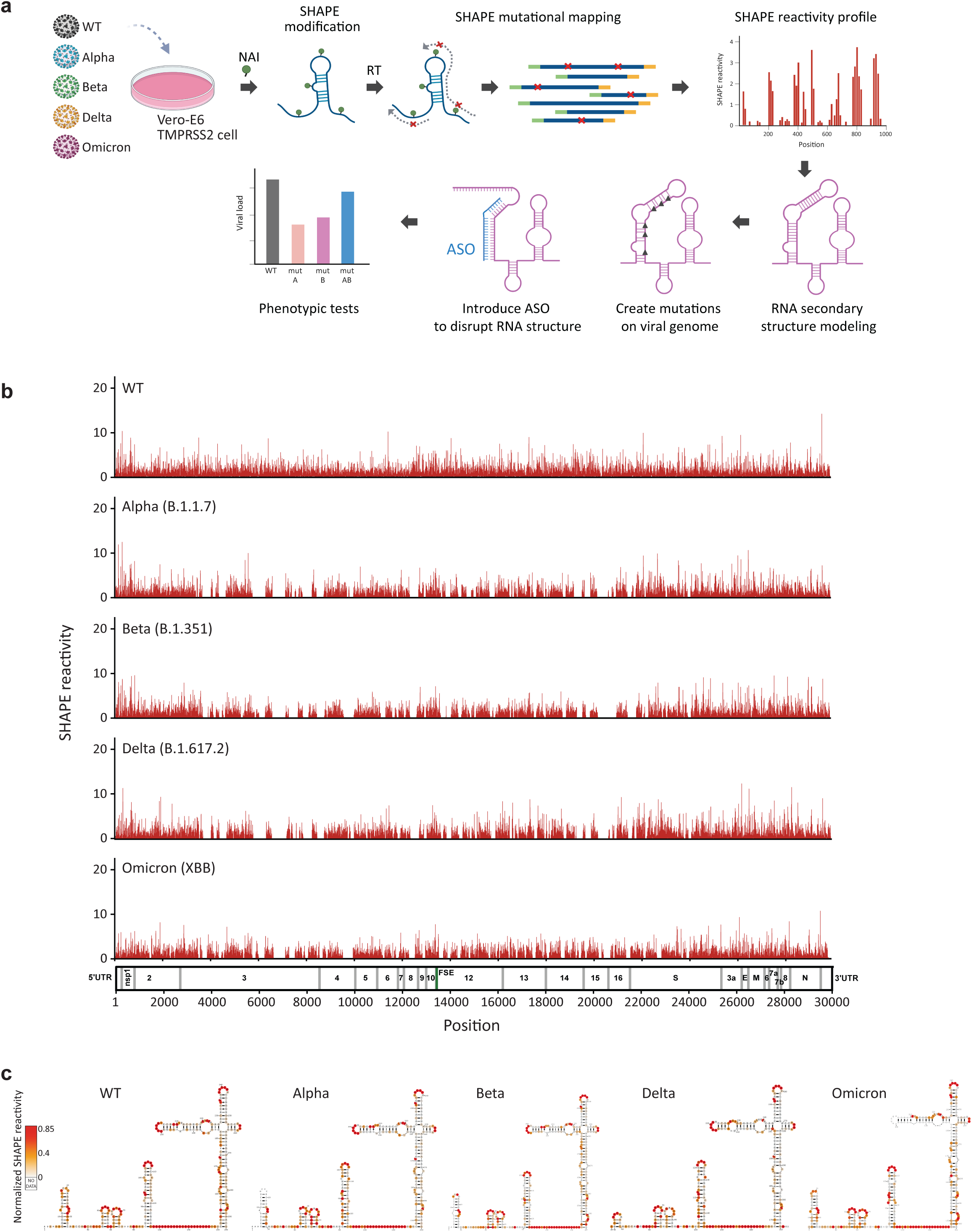
Genome-wide structure mapping of SARS-CoV-2 WT and VOCs in infected cells. **a** Schematic showing the workflow for SHAPE-MaP probing and identifying potential RNA structural elements with regulatory functions in five virus variants. Virus genomes were probed inside infected cells using NAI, a SHAPE-like chemical, which modifies single-stranded RNA bases along the viral genome. The SHAPE reactivity profiling data was then used to constrain computational models to derive more accurate structural predictions for SARS-CoV-2. **b** SHAPE-reactivity profiles for genomes of five SARS-CoV-2 variants. High reactivity indicates a higher probability of single-strandedness. **c** Structure models of 5’ UTR for five variants of SARS-CoV-2 created using the program RNAstructure with SHAPE-reactivities as constraints and visualized with VARNA. Source data are provided as a Source Data file.

We obtained a total of 689 million reads for at least 2 biological replicates (**Supp. Table 1**), of which 256 million (37.1%) were aligned to the SARS-CoV-2 variants. As expected, NAI-treated samples showed an increase in mutation rates as compared to untreated samples for all variants (**Supp. Figure 2a**). We also observed highly consistent SHAPE-reactivities between two replicates (*R*=0.86-0.93), indicating that our structure probing is robust (**Supp. Figure 2b**). Across the SARS-CoV-2 genomes, we were able to capture structural information for ∼84% of the bases (**Figure 1b, Supp. Figure 2c**), enabling us to identify RNA structures across the genomes of different variants. We confirmed that our SHAPE-reactivities recapitulate expected double and single-stranded RNA signals on the 5’ and 3’UTRs of the SARS-CoV-2 genome (**Figure 1c, Supp. Figure 3a,b**), and have high AUC-ROC against known SARS-CoV-2 structures (**Supp. Figure 3c**), indicating that our structure probing is accurate.

We had previously performed structure probing on a WT SARS-CoV-2 virus together with a 382-nucleotide deletion mutant (Δ382) to identify functional SARS-CoV-2 structures^15^. Now, probing RNA structures across five variants enabled us to identify a more conserved set of six low SHAPE, low Shannon (LSLS) regions along their genomes (**Figure 2a, Supp. Figure 4a**). Additionally, to identify structurally similar regions along the SARS-CoV-2 genomes, we performed Pearson correlation on sliding windows of SHAPE-reactivities along the different pairs of the genomes of SARS-CoV-2 variants (**Supp. Figure 4b**). We also used the reactivity along each virus genome to model local structures for the full genome and performed pair-wise local structure comparisons to identify the top 20 most structurally conserved elements across all the variants (**Figure 2a,b, Supp. Figure 5a, Supplementary Table 2**). These structures include known conserved features such as the 5’UTR, the FSE, and conserved LSLS regions in the Orf3a described above (**Figure 2a,b**). Using a set of 1581 sequences from the *Coronaviridae* family^12^, we observed that these 20 structures are supported by evolutionary covariation, indicating that they are likely to be functionally important (**Figure 2b**). To test whether these conserved structures are important for virus fitness, we designed an antisense oligonucleotide (ASO) against a region in Orf7a (residues 27695-27714 in WT RNA genome) that is highly base-paired and structurally conserved across the analysed viruses. We observed that transfecting cells with this ASO before WT and Omicron infection reduced the replication of both viruses, as compared to cells transfected with an ASO that targets a randomly selected region of the SARS-CoV-2 genome or with a non-targeting control ASO, indicating that this structure could be functionally important for virus replication (**Supp. Figure 6a,b,c**).

**Figure. 2.**
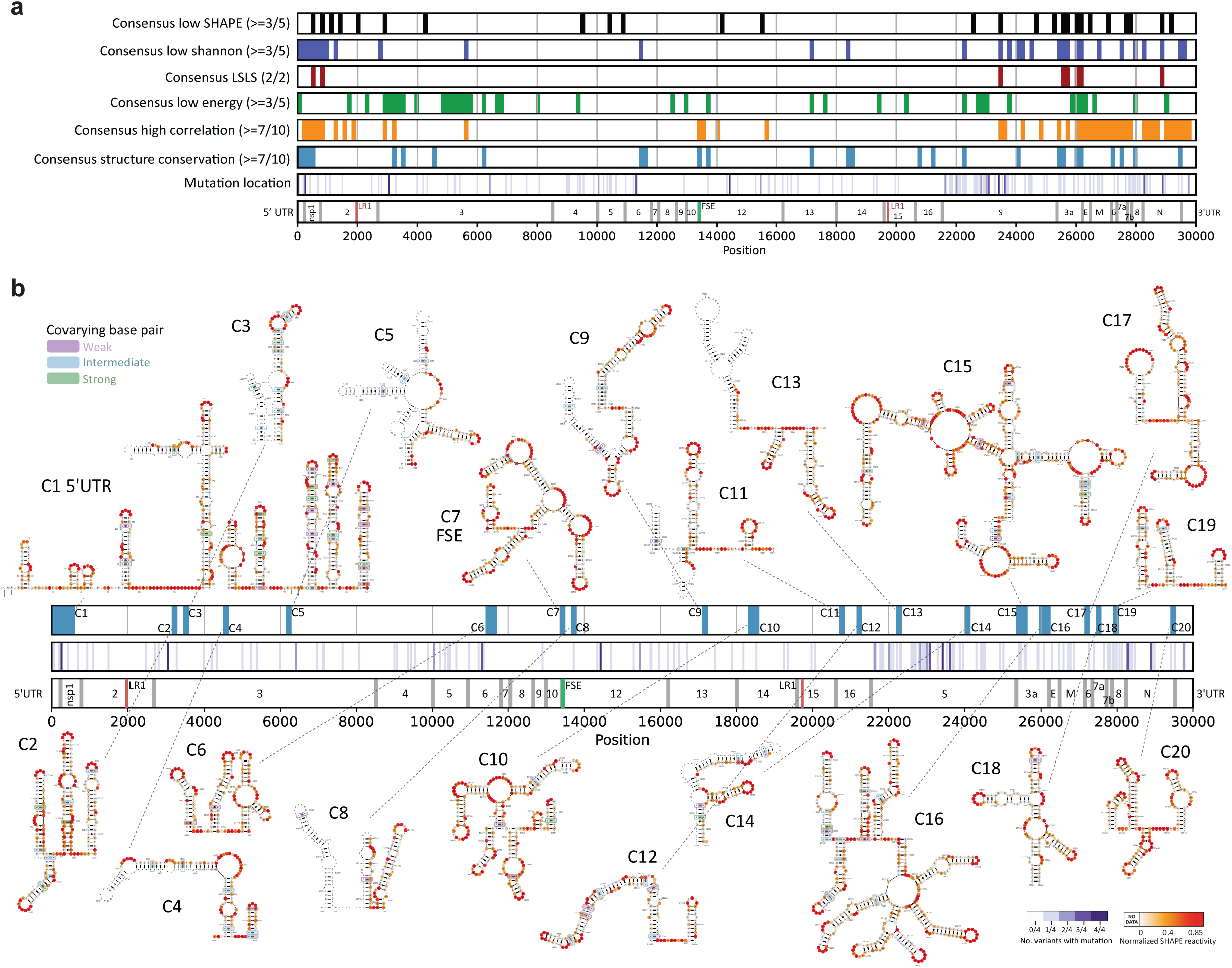
SARS-CoV-2 WT and VOCs share conserved RNA structures along the genomes. **a** Consensus regions (150 bases) with the lowest 20% SHAPE reactivity (black), Shannon entropy (blue), greatest difference (top 20%) between actual and shuffled energies (green), high SHAPE-MaP correlation (*R* ≥ 0.8, orange), and the top 20 structurally conserved regions (at least 85% identical base pairs, light blue) are shown along the SARS-CoV-2 genome map. **b** RNA structure models for the Omicron variant. The top 20 most conserved structures (light blue bars) were generated with the relevant RNAstructure program modules using SHAPE-MaP reactivity as a constraint and visualized with VARNA^46,50,51^. SHAPE-reactivities are mapped onto structure models as a low-to-high colour gradient. Base pair covariance was calculated via a protocol adapted from Sun, L. *et al*.^12^ and mapped to the corresponding pair with colour coding signifying strong (> 0.7), intermediate (0.5 - 0.7), and weak (0.4 - 0.5) covariance. The number of mutations (substitutions and indels) at a given genome position across the four VOCs (Alpha, Beta, Delta and Omicron) is displayed as purple gradients. The genome map shown for position reference is computed from a MSA of the WT SARS-CoV-2 and four VOCs. Source data are provided as a Source Data file.

### SNVs and RBPs impact virus structure along the virus genome

As the VOCs have different mutations along their genome (**Supp. Figure 1a,b,c**), we examined the impact of the SNVs on RNA structure differences between these viruses. Across WT and four VOCs, we observed 720 pair-wise sequence differences, with most of the bases changing from C to U (**Figure 3a**). Generally, RNA structures around SNVs were more variable than RNA structures around non-SNV bases (**Figure 3b, Supp. Figure 7a,b**). Out of 720 pair-wise mutations across the different viruses, we observed that 310 (∼43%) of them showed changes in RNA structures. Mutations that changed the original residue into purine bases (A/G) resulted in larger structural changes than SNVs that changed into pyrimidine bases (U/C, **Supp. Figure 8a**). As expected, bases that are mutated to A tend to become less paired, while bases that are mutated to G tend to become more paired (**Supp. Figure 8a**). As previously observed, RNA structure changes caused by SNVs are largely local and influence the reactivity of ∼10 bases up-and downstream of the SNV (**Figure 3c**). Scanning across the SARS-CoV-2 genome, we observed that one of the most structurally heterogeneous regions is associated with an SNV (C->U at base 7141 of the Delta variant, **Supp. Figure 5a**). Structure modelling of this region using SHAPE-reactivity as constraints showed that the region around position 7100 can form different structures in different variants (**Figure 3d**). Additionally, many SNVs do not result in local structure changes, as seen by the relatively structurally stable regions in the RNA region encoding for the S-protein, despite the large number of mutations present in the gene (**Figure 3e**). To determine whether other cellular factors such as RBPs can impact RNA structure changes in different SARS-CoV-2 variants, we identified 72 RBPs that are differentially expressed in Vero-E6 cells that are infected by different strains of SARS-CoV-2. These RBPs are experimentally identified to bind to SARS-CoV-2^12,19–23^, and are known to have binding motifs (**Figure 3f,g,h, Supp. Figure 9a,b,c**). While differences in RBP expression generally do not result in changes in RNA structures, we identified a few RBPs including HNRNPU and FXR1, whose expression was associated with the formation of different RNA structures in the SARS-CoV-2 genome (**Figure 3i,j, Supp. Figure 10a-c**). In particular, we observed that a highly variable region around position 11000 along the SARS-CoV-2 genome folds into different predicted structures in the presence of different levels of HNRNPU (**Figure 3j, Supp. Figure 10a**). To validate that HNRNPU binding indeed results in structure changes, we performed an in vitro assay by incubating a 500-base RNA fragment, centred around the HNRNPU binding site in the SARS-CoV-2 WT genome, in the presence or absence of the HNRNPU protein, and performing RNA structure probing and sequencing. Our SHAPE-MaP results show that adding HNRNPU protein results in an increased accessibility of a region upstream of the HNRNPU binding site (**Figure 3k**), similar to what was observed in infected cells, whereby high HNRNPU (in WT infected cells) results in an increased accessibility of a region upstream of the HNRNPU binding site, as compared to cells with low HNRNPU levels (In Beta infected cells, **Figure 3k, Supp. Figure 10a,b**). As HNRNPU is a novel RNA sensor involved in viral sensing, the different structures could be associated with differences in detecting different SARS-CoV-2 strains inside cells^24,25^.

**Figure. 3.**
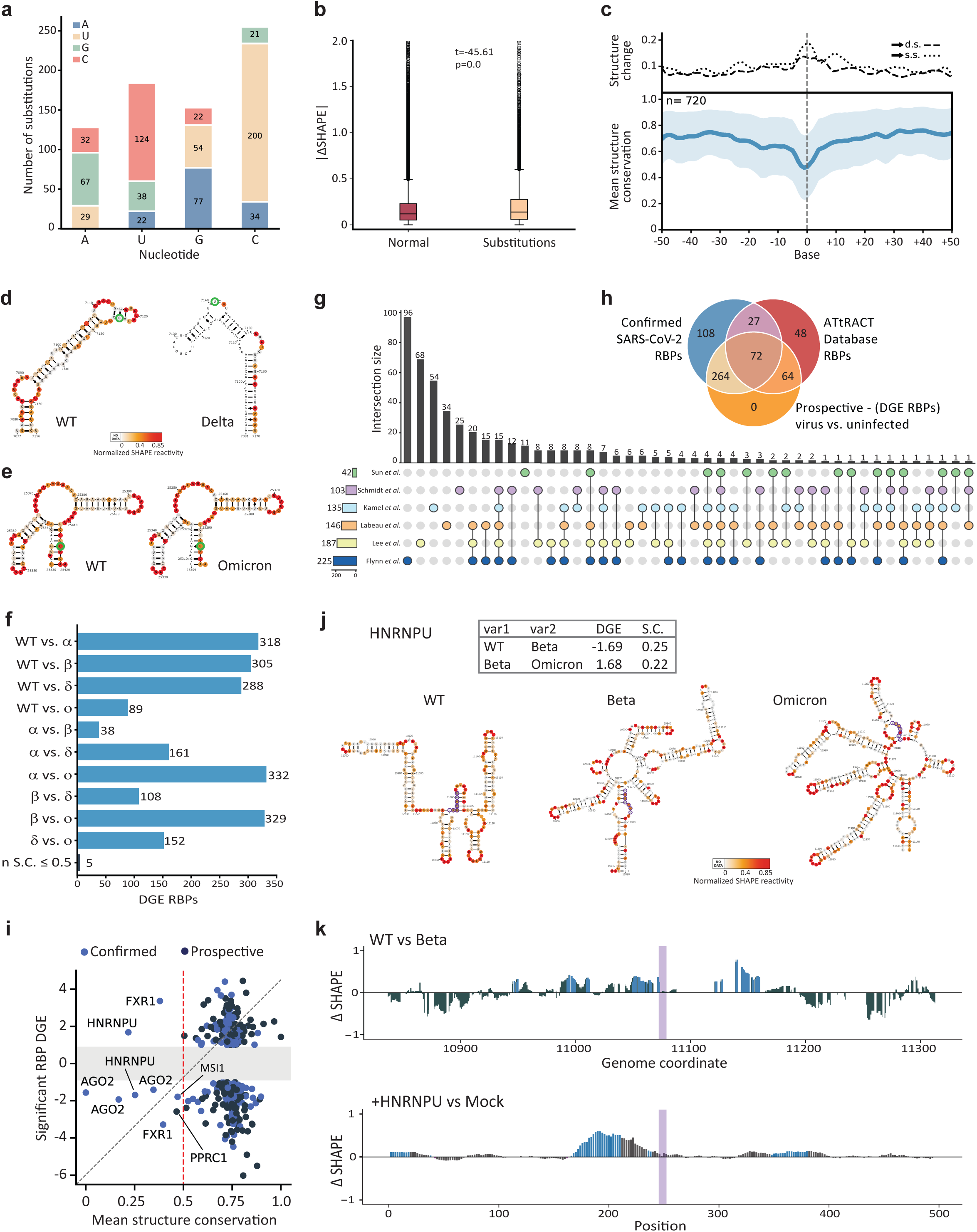
Nucleotide substitutions and RBPs binding cause structural changes along SARS-CoV-2 genomes. **a** Bar-chart exhibiting the counts and types of nucleotide substitutions found in all pairs of WT SARS-CoV-2 and VOC genomes. **b** Boxplot showing the distribution of change in SHAPE-reactivity in conserved versus variable bases along the SARS-CoV-2 variants. The box indicates the first and third quartiles, and the median is indicated as the line. **c** Top: Line plots showing the degree change as a base transitions to either double-stranded or single-stranded. Bottom: The line plot indicates the degree of structure similarity at each base, centred on the position of the base change. **d** RNA secondary structure models constrained by SHAPE reactivity in WT SARS-CoV-2 and Delta variant. The Delta variant has a single C to U substitution and shows an appreciable rearrangement in the base pairing configuration. **e** RNA secondary structure models constrained by SHAPE reactivity in WT SARS-CoV-2 and Omicron variant. The Omicron variant contains a single C to U substitution that does not result in a structural change. The loss of a GC Watson-Crick pair is compensated for by a GU wobble pair. **f** Bar graph illustrating the number of differentially expressed RBPs in each combination of WT and VOC pairs. **g** Upset plot detailing the overlap among six studies that define the SARS-CoV-2 RNA-bound proteome (RBPome). **h** Venn diagram depicting the intersection of several RBP sets from multiple databases. “Confirmed SARS-CoV-2 RBPs” are unique RBPs aggregated from literature sources, while “Prospective - (DGE RBPs)” are differentially expressed genes cross-referenced with “Confirmed” and “ATtRACT Database RBPs” lists. **i**. Scatterplot showing the correlation of RBP expression with the degree of structure similarity around the RBP binding motifs. RBPs with experimental evidence are coloured in light blue (Confirmed), while predicted RBPs from motif enrichments are coloured in black (Prospective). **j** Structure models of an RNA region that contains a HNRNPU motif in WT, Beta and Omicron. HNRNPU expression is different across cells infected with the different variants. While individual stem-loop structures are conserved, longer-range interactions are reorganized. **k** Top, DeltaSHAPE profile of HNRNPU binding region on SARS-CoV-2 genome comparing WT-infected to Beta-infected cells. Bottom, deltaSHAPE profile of an in vitro experiment using a 500-base RNA fragment that centred at the HNRNPU binding region, structure probed in the presence and absence of HNRNPU protein. Regions that are significantly less structured (positive change) are shown in light blue. The HNRNPU binding site is highlighted in a purple box. Source data are provided as a Source Data file.

### A long-range RNA-RNA interaction impacts virus fitness

As SARS-CoV-2 can form long-range RNA-RNA interactions within the genome that are hard to predict using SHAPE, we performed proximity ligation sequencing (SPLASH) to identify intra-and intermolecular RNA-RNA interactions in the cells infected with WT SARS-CoV-2 or the Omicron variant. This analysis identified 260 and 308 intramolecular interactions along the WT and Omicron genomes, respectively. We observed that 43-60% of the intramolecular RNA interactions in these two strains are long-range interactions that span more than 1 kb in distance (**Figure 4a,b, Supp. Figure 11a**). Out of all 260 SPLASH interactions in the WT strain, 172 (66%) are present in Omicron, including 73 out of 155 (47%) longer-range interactions (> 1 kb, **Figure 4c,d**). Both intra-and intermolecular RNA-RNA interactions are enriched in the structurally similar regions identified above (**Figure 4e,f**), although we do see an impact of SNV on intramolecular RNA interactions whereby only 54% of RNA-RNA interactions with SNVs (as compared to 66%) are conserved between WT and Omicron strains (**Figure 4g, Supp. Figure 11b**). SNVs preferentially disrupt longer-range (> 1 kb, 39% conservation between WT and Omicron) versus shorter-range (≤ 1 kb, 85% conservation between WT and Omicron) RNA-RNA interactions (**Figure 4h, Supp. Figure 11b**), suggesting that shorter-range RNA-RNA interactions are likely to be more conserved.

**Figure. 4.**
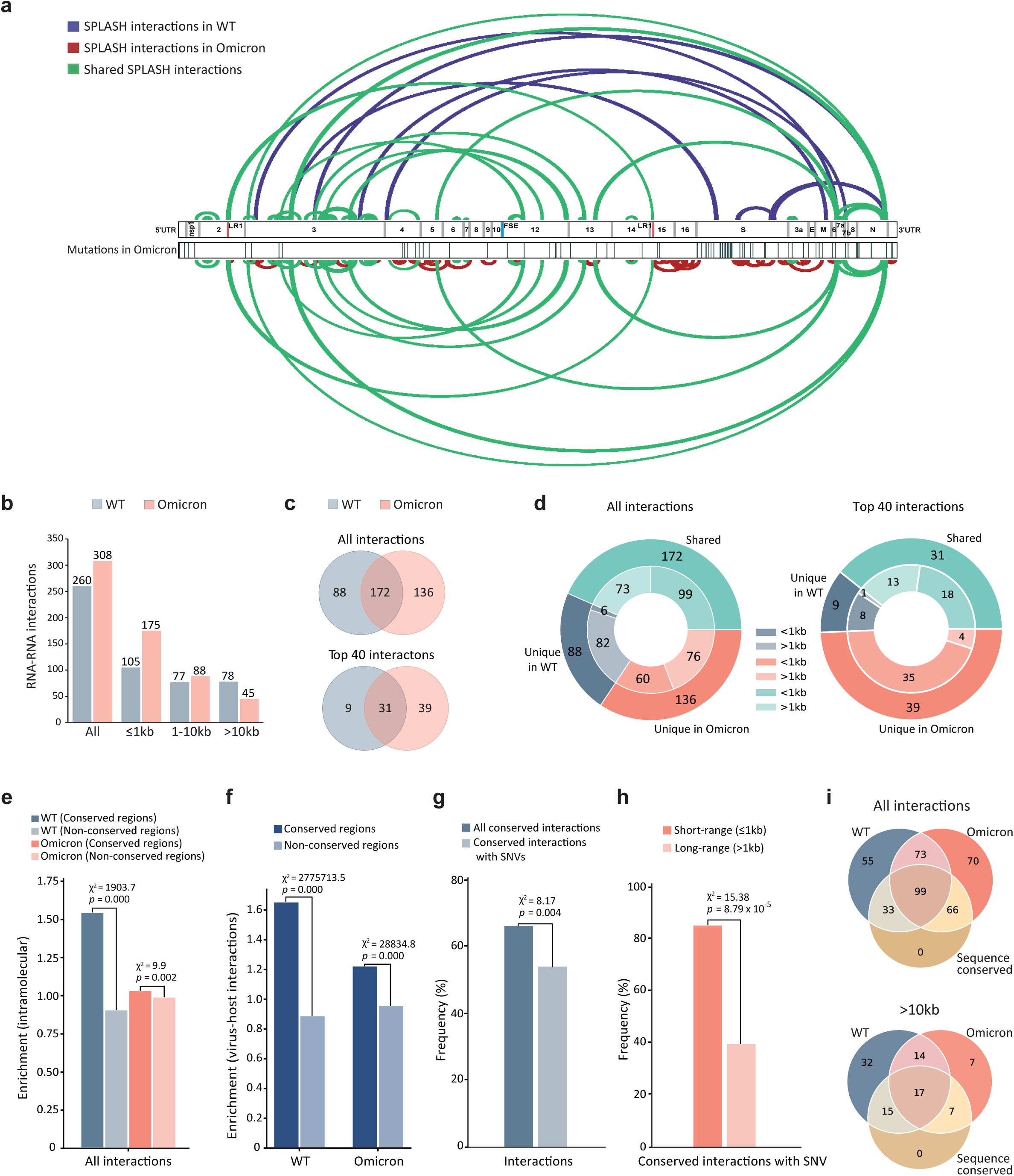
Conserved short-and long-range interactions in SARS-CoV-2 WT and Omicron genomes. **a** Arc plots showing top 40 pair-wise RNA-RNA interactions along the SARS-CoV-2 WT and Omicron genomes. Shared interactions between the WT and Omicron variants are highlighted in green. Unique interactions in WT (blue) and in Omicron (red) are plotted on the top and bottom panels, respectively. **b** Bar-chart showing the abundance of SPLASH interactions that span different lengths along the WT and Omicron genomes. Interactions over a distance longer than 1 kb are classified as long-range interactions which comprise 59.6% of all interactions in WT and 43.2% in Omicron, respectively. **c** Venn diagram showing the overlap of SPLASH interactions between SARS-CoV-2 WT (grey) and Omicron (pink) identified in infected Vero-E6/TMPRSS2 cells. **d** Multilevel doughnut chart showing unique or shared SPLASH interactions of SARS-CoV-2 WT and Omicron that span different distances along the genome. **e** Bar-chart showing interactions per base for short-(≤ 1 kb) and long-range (> 1 kb) interacting regions that fall within structurally conserved or non-conserved regions in WT and Omicron variants, respectively. **f** Bar-chart showing the number of RNA-RNA interactions between virus and host per base in structurally conserved and non-conserved regions. **g** Bar-chart showing the frequency of conserved interactions and conserved interactions with SNVs in WT and Omicron variants. **h** Bar-chart showing the frequency of conserved interactions with SNV that span short-range (≤ 1 kb) and long-range (> 1 kb) along the virus genome. P-values were calculated by Chi-square test. **i** Venn diagrams showing the intersection of RNA-RNA interactions that were experimentally identified from WT and Omicron-infected cells by SPLASH, as well as predicted interactions in all VOCs based on sequence conservation. Source data are provided as a Source Data file.

Out of a total of 260 RNA-RNA interactions detected, 78 interactions span longer than 10 kb in the WT strain. Sequence analysis revealed that from these ultra-long-range RNA-RNA interactions, 32 are conserved across all VOCs and 31 were experimentally confirmed to be conserved between WT SARS-CoV-2 and the Omicron variant (**Figure 4i, Supp. Figure 11c,d**). One of these conserved long-range RNA-RNA interactions, named LR1, brings bases around position 2000 (in the nsp2 encoding region), to interact with bases around position 19760 (in the nsp15 encoding region, **Figure 5a**). To explore the full ensemble of RNA structures that this interaction can form, we generated 1000 structures stochastically from the base-pairing probability of this region. We observed two main clusters, one corresponding to local structures (C1, C2, **Figure 5b**) and accounting for a third of all structures in the ensemble, and the rest corresponding to long-range interaction structures (C3, C4, **Figure 5b**).

**Figure. 5.**
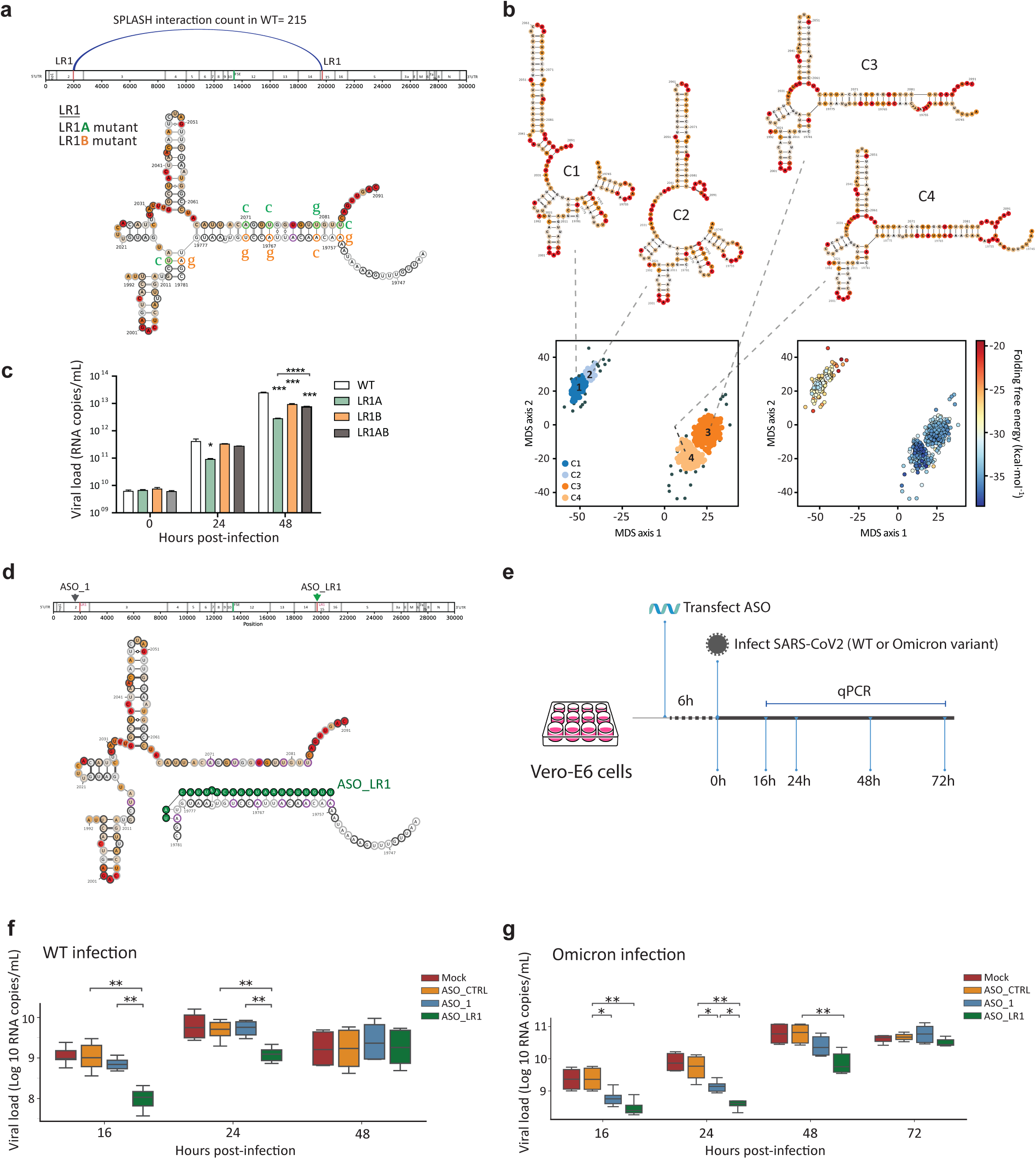
The functional importance of a long-range RNA-RNA interaction. **a** Top, Arc plot showing a conserved long-range RNA-RNA interaction (LR1) that spans 17 kb in distance along the SARS-CoV-2 genome. Bottom, structure model of the conserved LR1 interaction. The mutations for LR1A and LR1B mutant substitutions are indicated in green and orange, next to the original bases, respectively. Compensatory mutant LR1AB contains all the mutations for LR1A and LR1B to restore the proposed RNA structure. **b** Clustering analysis of the LR1 secondary structure ensemble reveals two populations of distinct structural morphology among the four clusters identified in the structural landscape. Long-range interaction structures in clusters C1 & C2 (dark and light blue dots) and clusters C3 & C4 (dark and light orange dots) are highly homologous, differing only by several base pair variations. Structure conformations (C3 & C4) that maximize long-range base pairing are more numerous and more stable than conformations that maximize local base pairing (C1 & C2). **c** Bar-chart showing the viral load of SARS-CoV-2-Wuhan-Hu1 (WT) and its mutants in the supernatants of infected Vero-E6 cells. Viral RNA was purified and quantified by RT-qPCR using an absolute quantification. Each column represents the mean ± SD of three technical replicates from one representative experiment. The same trends were observed in three independent experiments. **d** Diagram showing that antisense oligonucleotide (ASO_LR1) disrupts the original long-range base-pairing of LR1. **e** Workflow showing transfecting Vero-E6 cells with ASO, followed by SARS-CoV-2 WT or Omicron infection. Total RNA from infected cells was collected at the indicated timepoints and copy numbers of viral genomes were determined by RT-qPCR. **f,g** Box-plot showing the amount of SARS-CoV-2 WT and Omicron RNAs in cells that were transfected with different ASOs, at 16, 24 and 48 hpi. ASO_CTRL: ASO that does not target any region on the viral genome, ASO_1: ASO that targets a randomly selected region on the viral genome (position 1563-1582), ASO_LR1: ASO that targets LR1 to disrupt the long-range base-pairing structure. For data in **c**, *P*-values are calculated by Student’s t-test. For data in **f** and **g**, *P*-values were calculated by Mann-Whitney test. Source data are provided as a Source Data file.

To analyse if the long-range base-pairing is indeed important for SARS-CoV-2 fitness, we introduced silent mutations affecting one side of the interacting regions (LR1A), another side of the interacting regions (LR1B), or both sides (LR1AB) into the genome of the SARS-CoV-2-Wuhan-Hu1 strain (**Figure 5a**). Interestingly, we observed that mutating the A side resulted in lower copy numbers of the viruses being produced in infected cells at both 24 and 48 hpi, while mutating the B side resulted in a mild attenuation at 48 hpi. As the B side mutations introduced A to G substitutions, resulting in GU base-pairing, we expected the B mutations to be less disruptive, agreeing with our experimental data (**Figure 5a,c**). Introducing compensatory mutations on both A and B sides partially rescues the low copy number of the LR1A mutant, indicating that the base-pairing is important for virus fitness (**Figure 5c**). Interestingly, we observed that the mutation resulted in the reduction of virus titres only at late time points, 24 and 48 hpi. The mutant viruses generally displayed accelerated replication at earlier time points, suggesting that LR1 might influence later stages of the virus infection cycle, such as genome packaging (**Supp. Figure 12a-d**).

To confirm that this LR1 base-pairing is indeed important for virus fitness, we designed an ASO that is complementary to bases 19758-19779 to inhibit the original base-pairing of this region (**Figure 5d**). Transfecting cells with the ASO, followed by SARS-CoV-2 infection, reduced the replication of both the WT SARS-CoV-2 and Omicron variants while transfection of control ASOs did not (**Figure 5e,f,g**), supporting our hypothesis that this interaction is important for virus fitness across different strains. In vivo mouse experiments whereby mice that inhaled two different concentrations of LR1 ASO before being exposed to SARS-CoV-2 resulted in an attenuation of SARS-CoV-2 infection, including decreased weight loss and lower virus titre in mice (**Supp. Figure 12e-h**), suggesting that LR1 ASO could act as a prophylactic in mice.

To test which cellular factors could potentially bind to LR1 to regulate the growth of SARS-CoV-2, we generated a biotinylated version of a 158-base-long mini-construct that contains the LR1 base-pairing (mLR1) and incubated this RNA with cellular lysates (**Figure 6a**). RNA pull-down followed by mass spectrometry was used to identify proteins that could bind to mLR1. We identified ADAR1 as an RNA binding protein that interacts with mLR1 across three biological replicates (**Supp. Figure 13a, Supplementary Table 3**), with ADAR1 binding strongest to mLR1-WT, and less so to the mutants mLR1A and mLR1B RNA (**Supp. Figure 13b**). To confirm this interaction, we performed a Western blot experiment using lysates from Vero-E6. We observed that ADAR1 was pulled down with mLR1-WT and that this binding is significantly reduced in mutants mLR1A, mLR1B, and non-detectable for highly structured *Tetrahymena* RNA control (**Figure 6b**). We observed a similar result when the pull-down assays were performed using human lung cancer A549 cells (**Supp Figure 13c**). To confirm that ADAR1 binds to this long-range LR1 interaction region inside cells, we also performed two biological replicates of enhanced crosslinking and immunoprecipitation (eCLIP)^26^ using antibodies against ADAR1 (**Supp. Figure 13d**). We observed highly reproducible ADAR1 binding across the SARS-CoV-2 genome (**Supp. Figure 13e**), and that ADAR1 binding to the 5’ arm of LR1 in the SARS-CoV-2 genome was highly significant (**Supp. Figure 13f**). To test whether mLR1 indeed binds to ADAR1 directly, we performed an electrophoretic mobility shift assay (EMSA) using different amounts of ADAR1 protein. Again, we observed that WT mLR1 binds to ADAR1 the strongest, that this binding was partially disrupted by mutations introduced to the mLR1 (mLR1A or mLR1B) and is restored in the case of the mLR1AB compensatory mutant (**Figure 6c,d**). We also observed that binding between ADAR1 and highly structured *Tetrahymena* RNA was weak, suggesting that ADAR1 indeed specifically binds to LR1 and not to structured RNAs in general (**Figure 6c,d**).

**Figure 6.**
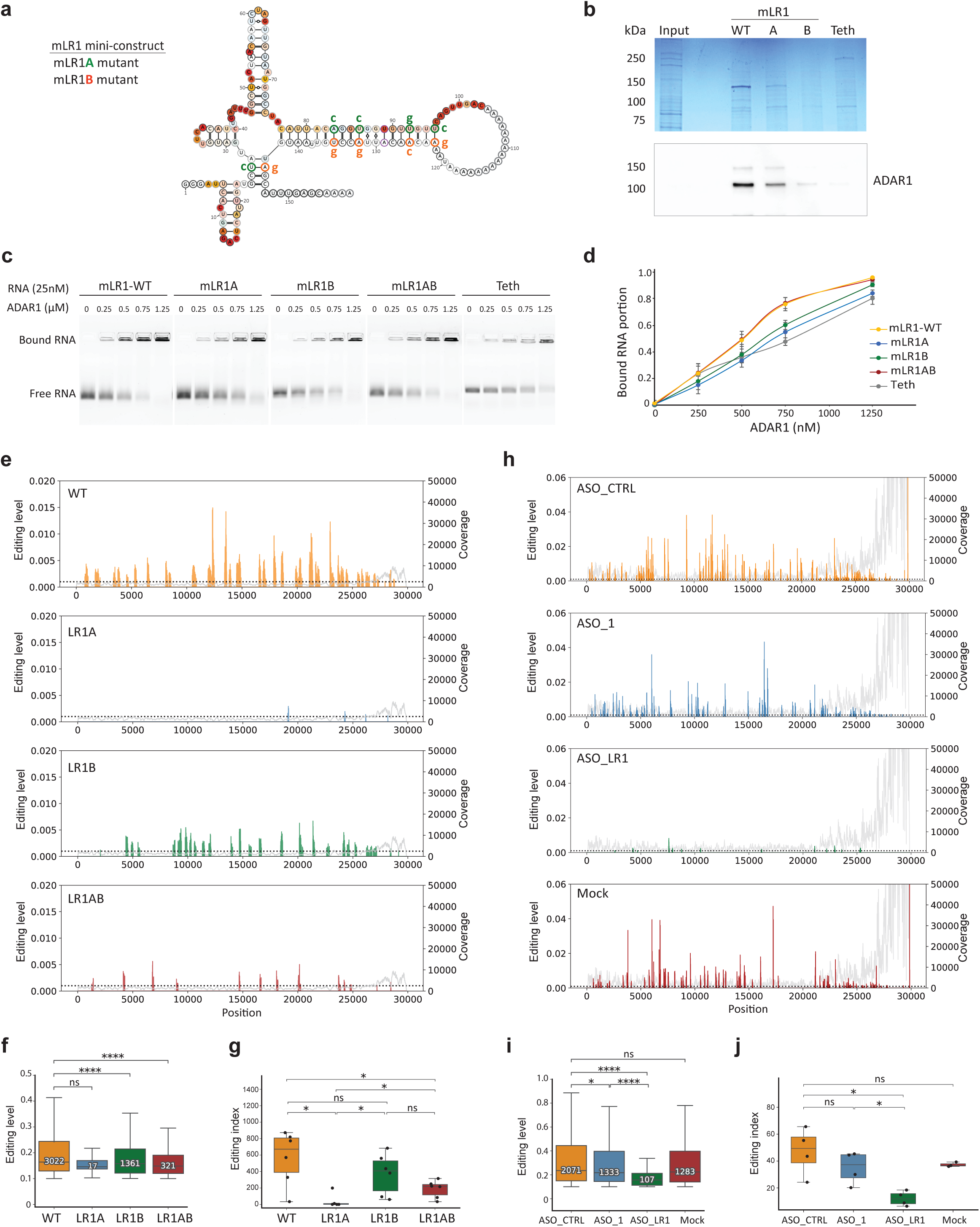
LR1 recruits ADAR1 to alter the editing level of the SARS-CoV-2 genome. **a** Structure model of a 158-base-long mini-construct that contains the long-range interaction LR1 base-pairing (mLR1). Mutations intended to disrupt the base-pairing from one side (mLR1A, green) or another side (mLR1B, orange) are indicated next to the original bases. Compensatory mutant mLR1AB contains both A and B mutations restoring the pairing of mLR1 to the original state. **b** Top, PAGE showing proteins that were immunoprecipitated using biotinylated mLR1-WT RNA and negative control RNAs mLR1A (lane A), mLR1B (lane B) and *Tetrahymena* ribozyme (Teth) using Coomassie blue staining. Input proteins from the whole cell lysate of Vero-E6 cells are also shown. Bottom, Western blot analysis of proteins from the pull-down assay using anti-ADAR1 antibody. **c** EMSA experiment showing the binding of different concentrations of ADAR1 (from 0 - 1.25 μM) with 25 nM of RNA from mini-construct mLR1-WT, mLR1A mutant, mLR1B mutant, mLR1AB compensatory mutant, and Teth as control. The samples are visualized using native agarose gel electrophoresis. **d** Quantification of the bound RNA fraction shown in **c** at the indicated ADAR1 concentrations. Three independent experiments were performed. Error bars are SD. **e** A-to-I RNA editing level along SARS-CoV-2 WT, LR1A, LR1B and LR1AB mutant genomes in infected Vero-E6 cells. The editing level was quantified by calculating the percentage of editing counts out of total reads at each position. Editing site calling was performed by subsampling each library to the same read depth for fair comparison. Analysis was performed in three independent experiments each containing two technical replicates **f** Boxplot showing the distribution of editing level of each edited site. The total number of edited sites across three biological replicates is indicated. *P*-value is calculated using the Wilcoxon Rank Sum Test. **g** Boxplot showing the distribution of the editing index on WT, LR1A, LR1B, and LR1AB mutants. The editing index was defined as the number of editing sites from edited viral reads divided by the total number of viral reads. **h** A-to-I RNA editing level along SARS-CoV-2 WT genome in mock-transfected cells (Mock), cells transfected using a random ASO (ASO_CTRL), an ASO that targets a different region (ASO_1, against 1563-1582), or an ASO designed against the LR1 region (ASO_LR1). **i** Boxplot showing the distribution of the editing level of each edited site along the SARS-CoV-2 WT genome in cells with different ASO treatment. Total number of edited sites across two biological replicates is indicated. **j** Boxplot showing the editing index in samples treated as described for panel **h**. *P*-values are calculated using the Wilcoxon Rank Sum Test. Source data are provided as a Source Data file.

ADAR1 has been shown to edit the genome of SARS-CoV-2. Additionally, COVID-19 patients have also been observed to have increased A-to-I editing levels along host transcripts^27,28^. To confirm that ADAR1 binding results in changes in RNA editing on the SARS-CoV-2 genome, we performed RNA sequencing analysis on Vero-E6 cells that are infected by SARS-CoV-2-Wuhan-Hu1 (WT), LR1A mutant, LR1B mutant, and LR1AB compensatory mutant viruses. Across three biological replicates, we identified a total of 3022 A-to-I editing sites in the WT, 17 in the LR1A mutant, 1361 in the LR1B mutant, and 321 in the LR1AB mutant genomes (**Figure 6e**). This suggests that A-to-I editing is severely repressed when LR1 is mutated and that restoring the base-pairing in the LR1AB mutant partially restores the number of editing sites on the genome. In addition to calculating the number of editing sites, we also calculated an editing index for each genome, which is the normalized number of editing sites that reflects the editing status of each sample. We observed that the LR1A mutant showed significantly lower editing index as compared to the WT, LR1B, and LR1AB mutants (**Figure 6g**), indicating that it has lower editing levels. We confirmed our observation by calculating A-to-I editing upon ASO disruption of the LR1 pairing. Treating the infected Vero-E6 cells with a control ASO that does not target the SARS-CoV-2 genome (2071 A-to-I sites) and with an ASO that targets another region of the SARS-CoV-2 genome (1333 A-to-I sites) resulted in high levels of A-to-I editing along the virus genome as was also observed in untreated cells (1283 A-to-I sites). However, transfecting cells with ASO against LR1 prior to virus infection resulted in a ten-fold decrease of A-to-I editing sites (107 A-to-I sites), confirming that disrupting LR1 impacts A-to-I editing along the virus genome (**Figure 6h**). Consistently, the editing index of ASO_LR1-treated cells was also significantly reduced as compared to control ASO-treated cells as well as untreated cells (**Figure 6j**). In addition to quantifying the number of editing sites, we also quantified the percentage of editing at each edited site (editing level). We observed a decrease in editing levels upon mutating LR1 which was, again, partially restored by compensatory mutations (**Figure 6f**). The same effect was observed when transfecting cells with the ASO designed against LR1, and not with the ASO against another part of the virus genome (**Figure 6i**), confirming that LR1 binding to ADAR1 results in editing of the SARS-CoV-2 genome.

To determine whether this decrease in A-to-I editing in the LR1 mutants is due to decreased levels of ADAR1 expression in the infected cells, we performed RT-qPCR analysis to determine the amount of ADAR1 mRNA expression in SARS-CoV-2-Wuhan-Hu1 (WT), LR1A, LR1B, and LR1AB mutants infected cells. Interestingly, we observed a similar level of ADAR1 expression in all the infected cells (**Supp. Figure 14a**), indicating that the low RNA editing levels in LR1 mutants were not due to changes in ADAR1 levels, but likely due to differential recruitment of ADAR1. As ADAR1 has been shown to bind to SARS-CoV-2 nucleocapsid (N) protein to interact with the SARS-CoV-2 genome^29^, we also tested for the ability of mLR1 WT and mutant RNAs to bind to N protein by using the biotinylated mLR1 mini-constructs. Interestingly, we observed that RNA containing WT mLR1 binds strongly to the N protein, the binding is disrupted in mLR1 mutants (mLR1A, mLR1B), and partially restored in mLR1AB compensatory mutant (**Supp. Figure 14b,c**), suggesting that both ADAR1 and N protein co-bind to the SARS-CoV-2 genome. We also observed that WT and mLR1 mutants do not bind to SARS-CoV-2 envelope (E) protein, suggesting that mLR1 binding to ADAR1 and N proteins is specific (**Supp. Figure 14d**). This is consistent with our observation above that the LR1 mutation impacts late stages of the virus infection and could be impacting genome encapsulation of the SARS-CoV-2.

## DISCUSSION

Numerous mutations have emerged in the SARS-CoV-2 genome during the years of the COVID-19 pandemic. These changes have increased transmission of SARS-CoV-2 as well as allowed escape from immune responses. However, how they impact RNA structures in SARS-CoV-2 RNA is still understudied. Additionally, identifying ultra-long-range RNA-RNA base pairing and understanding their functions for RNA viruses remains a challenge in the field due to the extensive amount of search space that these interactions can form. Here, we utilized high-throughput structure mapping to map RNA structures across five different variants of SARS-CoV-2 to identify structurally similar RNA regions. We also used proximity ligation sequencing to identify shared long-range RNA-RNA interactions in the WT SARS-CoV-2 and Omicron variant and observed that SNVs preferentially impact long-range RNA structures. We identified 78 long-range RNA-RNA interactions that span a distance of more than 10 kb, out of them 31 were present in both the WT SARS-CoV-2 and Omicron variants. Interestingly, one of these interactions spans 17 kb in distance, can result in two alternative structures, and is important for virus fitness. Our experiments show that this long-range RNA element binds directly to ADAR1 and is associated with increased RNA editing along the SARS-CoV-2 genome.

A-to-I editing has been associated with SARS-CoV-2 infection^27,30^, however how it impacts viral infection still remains to be fully understood. Large-scale proteome screens have identified ADAR1 as an interaction partner with the SARS-CoV-2 genome^20,22^. Previous literature has also shown that the SARS-CoV-2 genome undergoes A-to-I editing, and that the transcriptome of SARS-CoV-2-infected cells shows elevated A-to-I editing^28,30,31^. Here, we identified an RNA element that could potentially recruit ADAR1 for RNA editing of the SARS-CoV-2 genome. As SARS-CoV-2 is sensitive to the interferon response and A-to-I editing of the virus genome could dampen the interferon response upon infection, we hypothesize that the editing of the SARS-CoV-2 genome could impact cellular sensing of the viral RNA, enabling the virus to replicate better inside cells. However, ADAR1 alone might not be s sufficient for this effect to occur. ADAR1 is shown to work together with other proteins such as DHX9, which is another top hit in our LR1 mass spectrometry data, to impact RNA sensing, complicating our understanding of how SARS-CoV-2 can be recognized inside cells^32^. Additionally, as disrupting the LR1 long-range interaction impacts virus fitness in the late stages of its lifecycle, and we observed binding of nucleocapsid protein with LR1, we further hypothesize that LR1 might be involved in packaging of the SARS-CoV-2 genomes through its interaction with the nucleocapsid protein and ADAR1. Our study highlights the complexity of SARS-CoV-2 RNA structures and demonstrates that ultra-long-range RNA-RNA interactions within the RNA genome can be functionally important by recruiting host proteins affecting its infectivity.

## METHODS AND MATERIALS

### Cells and viruses

African green monkey Vero-E6 cells (ATCC CRL-1586) and Vero-E6 cells constitutively expressing human TMPRSS2 (Vero-E6/TMPRSS2; BPS Bioscience #78081) were used in this study. Vero-E6 cells were cultured in Dulbecco’s Modified Eagle Medium (DMEM; Cytiva, Utah, USA) supplemented with 10% (v/v) heat-inactivated fetal bovine serum (FBS; Capricorn Scientific, Ebsdorfergrund, Germany). Vero-E6/TMPRSS2 cells were grown in DMEM supplemented with 10% (v/v) heat-inactivated FBS, 1% (v/v) antibiotic-antimycotic (Gibco, Carlsbad, CA, USA) and 3 µg/mL of puromycin (InvivoGen, San Diego, CA, USA). Both cell lines were maintained at 37°C in a humidified 5% CO_2_ incubator.

The five SARS-CoV-2 variants used in this study were: SARS-CoV-2 wildtype (WT; Clade L, lineage B, hCoV-19/Singapore/1017/2020, EPI_ISL_574502), Alpha (lineage B.1.1.7, EPI_ISL_754083), Beta (lineage B.1.351.3, EPI_ISL_1173248), Delta (lineage B.1.617.2, EPI_ISL_2621925) and Omicron XBB (EPI_ISL_14917728). The viruses were isolated from the nasopharyngeal swabs of patients with COVID-19 confirmed by RT-qPCR. The virus isolates were propagated in Vero-E6 cells with no more than 3 passages prior to downstream experiments. All virus work was performed in the Biosafety Level 3 (BSL-3) laboratory and all protocols were approved by the BSL-3 Biosafety Committee and Institutional Biosafety Committee of the National University of Singapore.

### SHAPE-MaP structure probing of five SARS-CoV-2 viruses in Vero-E6 cells

Vero-E6/TMPRSS2 cells were seeded in T75 flasks (Corning, NY, USA) at a seeding density of 2.5 x 10^6^ cells/flask and incubated overnight. Cells were infected with SARS-CoV-2 WT, Alpha, Beta, Delta or Omicron XBB variants at a multiplicity of infection (MOI) of 0.01. At 1 hpi, inoculum was replaced with DMEM supplemented with 2% FBS, followed by incubation at 37°C. At 48 hpi cells were washed once with PBS, trypsinized and collected by centrifugation at 300 x g for 5 min. The cell pellet was then resuspended in PBS and divided into three experimental samples: (i) 1:20 volume of 1 M 2-methylnicotinic acid imidazolide (NAI; final concentration of 50 mM) was added and cells were incubated at 37°C for 15 min; (ii) 1:20 volume of dimethyl sulfoxide (DMSO) was added and cells were incubated at 37°C for 15 min; and (iii) cells were left without any treatment. Total RNA was extracted using RNeasy Mini Kit (Qiagen, Valencia, CA, USA) according to the manufacturer’s instructions. The harvested RNA was heat-inactivated at 65°C for 10 min before transferring it to BSL-2 laboratory for construction of cDNA libraries compatible for Illumina sequencing following the SHAPE-MaP protocol^33^.

### Interactome mapping of SARS-CoV-2 viruses in Vero-E6 cells by SPLASH

Vero-E6/TMPRSS2 cells were seeded in 15 cm cell-culture dish (Corning, NY, USA) at a seeding density of 2.5 x 10^5^ cells/mL and incubated overnight. Cells were then infected with SARS-CoV-2 WT or Omicron XBB at a MOI of 0.01. At 48 hpi, cells were washed once with PBS and incubated with biotinylated psoralen (BioPso; final concentration of 200 µM) and 0.01% digitonin in PBS at 37°C for 5 min. The BioPso-treated cells were then irradiated at 365 nm on ice for 20 min using a CL-1000 ultraviolet crosslinker (UVP Inc.). Total cellular RNA was extracted using TRIzol LS reagent (Invitrogen, Carlsbad, CA, USA) according to the manufacturer’s recommendations, followed by heat-inactivation at 65°C for 10 min before transferring it to BSL-2 laboratory for preparation of SPLASH libraries following the published protocol^34^.

### Construction of SARS-CoV-2 mutants

The SARS-CoV-2 mutants (LR1A, LR1B and LR1AB) were assembled using conventional restriction enzyme-based cloning, following previously described methods^35^. Briefly, sub-fragments containing mutations (Genscript) were introduced into pCCI-4K-SARS-CoV-2-Wuhan-Hu1 (MT926410) infectious cDNA (icDNA) plasmid using corresponding restriction enzymes. The right constructs were confirmed with Sanger sequencing. Five microgram of plasmid DNA (SARS-CoV icDNA and its mutants) were transfected into BHK21-ACE2-N cells (a gift from Suzannah J. Rihn) grown in T25 flasks using lipofectamine LTX reagent (Thermo Fisher Scientific) following manufacturer’s instructions. After 4-5 days post-transfection, the supernatant was transferred to Vero-E6 cells in T25 flasks. Rescued viruses (P1 stocks) were collected 3-4 days post-infection and titred using focus-forming assay. All work with recombinant viruses was performed in the BSL-3 laboratory of University of Tartu.

### Analysis of viral RNA kinetics by RT-qPCR

Vero-E6 cells were seeded in 6-well plates at a density of 4 x 10^5^ cells per well one day before infection. The cells were infected with SARS-CoV-2-Wuhan-Hu1 (WT) or its mutants at a MOI of 0.001. Experiment was performed in triplicates and 0.5 mL inoculum (DMEM + 0.2% BSA) was used per well. At 1 hpi, the inoculum was replaced with 2 mL viral growth media and cells were cultured for 30 min to collect the 0 hpi sample. Samples for isolation of extracellular and intracellular viral RNAs were collected at 0, 6, 12, 24 and 48 hpi for RT-qPCR. For each timepoint, the media was collected, and cells were washed with PBS, then trypsined, and collected in 200 μL of 1x shield buffer supplemented with 10 μL protease K (R2132, Zymo Research Quick-RNA MagBead). The cell lysates were frozen at-80°C for later RNA purification. Extracellular viral RNA was isolated using the MagMAX™ Viral/Pathogen Nucleic Acid Isolation Kit (A42352, Thermo Fisher Scientific) according to the manufacturer’s manual, utilizing the KingFisher™ Flex in a 96 deep well format. The purified extracellular viral RNA was then analysed using RT-qPCR by LightMix Modular SARS-CoV-2 (COVID-19) RdRp 530 (Roche), as described previously^36^. For analysis of intracellular viral RNAs, total cellular RNAs were purified using KingFisher Flex with the R2132 cells Quick RNA Magbead KingFisher Flex protocol. The cDNA of viral positive, negative and sub-genomic RNAs were synthesized using strand-specific primer according to the First strand cDNA synthesis kit manual (K1612, Thermo Fisher Scientific). PowerTrack SYBR Green Master Mix (A46109, Thermo Scientific) was used for qPCR analysis. The qPCR primers used for positive-and negative-strand RNA were SARS2-RdRp-F and SARS2-RdRp-R. The qPCR primers used for sub-genomic RNA were SARS2-N-F and SARS2-N-R, while the qPCR primers used to detect mRNA of GAPDH control were GAPDH-F and GAPDH-R. The obtained data was analysed using 2^-ΔΔCt^ method. The sequences of primers used for cDNA synthesis and for qPCR are listed in Supplementary Table 4.

### SARS-CoV-2 viral fitness assay

Vero-E6 cells seeded at a density of 2 x 10^5^ cells per well in 24-well plates one day before transfection were grown in high glucose DMEM media supplemented with 5% FBS. The next day, cells were transfected with 100 nM ASO using Lipofectamine RNAiMAX reagent (Thermo Scientific). Six hours after ASO transfection, cells were infected with WT SARS-CoV-2 or Omicron XBB variant at a MOI of 0.01. Infected cells were collected at 16, 24, 48 and 72 hpi, total RNAs were extracted by Qiagen Viral RNA Mini Kit. Viral RNA copy numbers were examined by RT-qPCR using Invitrogen SuperScript III Platinum One-Step RT-qPCR Kit according to the manufacturer’s instructions. The ASO sequences are listed in Supplementary Table 4.

### RNA pull-down assay

In vitro transcribed RNAs (mLR1 mini-construct and highly structured *Tetrahymena* RNA used as negative control) were biotinylated using Pierce RNA 3’ End Biotinylation Kit. RNAs were heated, put on ice for 2 min and RNA structure buffer (50 mM Tris pH 7.4, 10 mM MgCl_2_, 150 mM NaCl) was added. To allow proper folding the temperature was gradually increased from 0°C to 37°C. Vero-E6 or human lung cancer cells (A549) were trypsinized and collected by centrifugation at 300 × g for 5 min. Cells were lysed in Lysis/Binding buffer (50 mM Tris HCl, pH 7.5, 100 mM KCl, 5 mM MgCl_2_, 10% glycerol, 0.2% NP-40, 0.5 mM DTT, 1x protease inhibitor cocktail (Calbiochem), 50 U/mL SUPERase·In™ RNase Inhibitor) at 4°C for 10 min. Lysates were centrifugated at 1300 × g for 5 min to remove cell debris and obtained supernatants were collected in new tubes. Twenty picomole of folded RNA was then added to the cell lysates and incubated at room temperature for 2 h with rotation. Next, twenty microliters washed Dynabeads MyOne Streptavidin C1 beads (Invitrogen) were added followed by further incubation at room temperature for 1 h. Beads were washed 5 times and enriched proteins were eluted in 1x protein loading buffer by heating at 95°C for 5 min. Retrieved proteins were resolved using SDS-PAGE for mass spectrometry and Western blot analyses. For mass spectrometry analysis, pull-down proteins were resolved using SDS-PAGE followed by bands visualization using PageBlue protein staining solution (Thermo Scientific). Subsequently, areas of interest were excised from Coomassie-stained gel and subjected to in-gel digestion with Trypsin, following general in-gel protocol^37^. Following peptide extraction, samples were vacuum dried, resuspended in solvent A (0.1% formic acid in water) and equal amount of sample was loaded on the C-18 Evotip (Evosep) according to manufacturer instructions. Samples were analyzed using Evosep liquid chromatography system (Evosep) coupled with a timsTOF-SCP (Bruker), with a 15 cm x 75 μm column Aurora Elite (IonOptics), in DIA-PASEF mode. Each sample was separated by SPD40 method using 0.1% formic acid in water and 99.9% acetonitrile with 0.1% formic acid. DIA-PASEF data was searched using DIA-NN 2.0 software^38^ using in-silico library created from Uniprot fasta for African green monkey (*Chlorocebus sabaeus*). Western blot analysis was performed according to standard procedures using a 1:1000 dilution of anti-ADAR1 (E6X9R) XP(R) rabbit monoclonal antibody (Cell Signalling Technology) and a 1:5000 dilution of secondary anti-rabbit antibody. The sequences of mLR1 mini-constructs and *Tetrahymena* are listed in Supplementary Table 4.

### ADAR1 eCLIP experiment

The ADAR1 eCLIP experiment was performed as published protocol^26^. Vero-E6 cells were seeded in 10 cm cell-culture dish (Corning, NY, USA) at a seeding density of 2.5 x 10^5^ cells/mL and incubated overnight. Cells were then infected with SARS-CoV-2 WT at a MOI of 0.01. At 48 hpi, cells were washed once with pre-chilled PBS, UV crosslinked at 400 mJ/cm^2^ on ice at 254 nm radiation. Crosslinked cells were then scraped and collected by centrifugation at 300 × g for 5 min at 4°C. Cell pellets were lysed using iCLIP buffer (50 mM Tris-HCl pH 7.4, 100 mM NaCl, 1% NP-40, 0.1% SDS, 0.5% sodium deoxycholate), lysates were sonicated and digested with DNase and diluted RNase I. Cell lysates were immunoprecipitated using 10 µg of ADAR1(E6X9R) XP(R) rabbit monoclonal antibody (Cell Signalling Technology) overnight at 4°C. Two percent volume of each lysate sample was collected for preparation of a parallel Input library. The remaining cell lysates were processed as the immunoprecipitated (IP) samples and washed by high salt buffer (50 mM Tris-HCl pH 7.4, 1 M NaCl, 1 mM EDTA, 1% NP-40, 0.1% SDS, 0.5% sodium deoxycholate) to remove unspecific protein-protein interactions. Bound RNA fragments were dephosphorylated and ligated to an RNA adaptor at 3’ end. Protein-RNA complexes from Input and ADAR1-IP samples were run on 4-12% BisTris SDS-PAGE and transferred to nitrocellulose membrane. Membrane regions comprising the protein sizes to 110 kDa and above were excised, and RNA was released from the complexes with proteinase K. Input samples were dephosphorylated and 3’-end ligated to an RNA adaptor. All samples were reverse transcribed and cDNAs were 5’-end ligated to a DNA adaptor. Library was constructed and sequencing was performed using NovaSeq X Plus PE150.

### Electrophoretic mobility shift assay (EMSA)

The assay was performed using agarose gel as previously described^39^ with some modifications. Twenty five nanomole of folded RNA was incubated with increasing concentration (0 to 1.25 μM) of ADAR1 protein (TP319761M, Origene), SARS-CoV-2 nucleocapsid (N) protein (Z03488, GenScript), or SARS-CoV-2 envelope (E) protein (RP-87682, Thermo Scientific) in binding buffer (20 mM Tris buffer pH 7.5, 2 mM DTT, 2 mM MgCl_2_, 15 mM NaCl) for 1 h at 4°C. Sample was then mixed with 6x DNA loading buffer (R0611, Thermo Scientific) and loaded on 1% agarose gel with recommended concentration of GelRed in 0.5x Tris/Boric acid/EDTA (TBE) buffer. The electrophoresis was performed at 10V/cm for 20 min at room temperature, gels were imaged using a ChemicDoc (Bio-rad).

### Mice experiments

Animal experiments were conducted in a pathogen-free Animal Biosafety Level 2 (ABSL-2) facility in accordance with the National University of Singapore (NUS) Institutional Animal Care and Use Committee (IACUC) (Protocol No. R24i-0008) and the NUS Institutional Biosafety Committee (IBC) approved SOPs. Eight-week-old female K18-hACE2 transgenic mice (InVivos Ptd Ltd., Singapore) were used in this study. Mice were randomized and assigned to control and treatment groups. Studies were conducted in a blinded manner. The mice were acclimatized in the ABSL-2 facility for 72 h prior to the start of the experiment with 12 h light/dark cycles and controlled temperature (23 ± 2°C) and humidity (50 ± 10%). K18-hACE2 transgenic mice were subjected to a four-day pre-treatment regimen with saline, low dose (400 µg) or high dose (1000 µg) of ASO LR1 via intranasal route once daily. The viral challenge was conducted through intranasal delivery in 25 µL (split across both nostrils) of approximately 1 x 10^3^ PFU of ancestral SARS-CoV-2 WT variant. Body weight and physiological conditions were monitored daily throughout the experiment or until the humane endpoint was reached. Mice were sacrificed 4 days post-infection (dpi), with lung tissues harvested. Each organ was halved for the plaque assay (right lobes) and histopathological analysis (left lobe). To assess the viral load by plaque assay, lung tissues were homogenized with 500 µL of DMEM (Cytiva) supplemented with antibiotic and antimycotic (Gibco) and titrated in A549-hACE2 cells. The viral supernatants were serially diluted in 10-fold increments, and 100 µL of each serially diluted supernatant was added to the confluent A549-hACE2 cells into 24-well plates. After 1 h of incubation for viral adsorption, the inoculum was removed and washed once with PBS. About 1.2% microcrystalline cellulose-DMEM supplemented with antibiotic and antimycotic overlay media was added to each well and incubated at 37°C, 5% CO_2_ for 72 h. The cells were then fixed in 10% formalin and counterstained with crystal violet. The number of plaques were determined, and the virus titre of individual sample was expressed in logarithm of plaque forming unit (PFU) per organ. For histopathological analysis, left lung lobes were fixed in 3.7% formaldehyde overnight. Tissues were routinely processed, embedded in paraffin blocks (Leica Surgipath Paraplast), sectioned at 4 µm thickness, and stained with haematoxylin and eosin (Thermo Scientific) following standard histological procedures. For immunohistochemistry (IHC), the lung sections were deparaffinized and rehydrated, followed by heat-mediated antigen retrieval, quenching of endogenous peroxidases and protein blocking. Sections were then covered with rabbit anti-SARS-CoV-2 nucleocapsid protein mAb (Abcam, 1:1000 dilution) for 1 h at room temperature. Subsequently, the sections were incubated with rabbit-specific HRP polymer secondary antibody (Abcam, no dilution), visualized using chromogenic substrate DAB solution, and counterstained with haematoxylin.

### SHAPE-MaP analysis

Sequencing reads obtained from two replicates of SHAPE experiments were aligned with the respective sequences for the SARS-CoV-2 viruses (WT: EPI_ISL_574502, Alpha: EPI_ISL_754083, Beta: EPI_ISL_1173248, Delta: EPI_ISL_2621925, Omicron: EPI_ISL_14917728) and SHAPE values for each position were calculated using Shapemapper-2.1.5 with Bowtie-2.4.2 for reading alignment^40^. Read depths obtained in the sequencing experiments allowed for conclusive determination of SHAPE reactivities at approximately 74-99% of positions. A reference alignment of WT, Alpha, Beta, Delta and Omicron sequences was generated using mafft-7.520 with the L-INS-I strategy. Subsequently, the local correlation of SHAPE reactivity between replicates was calculated using Pearson correlation. As Pearson correlation between replicates for each data set was ≥0.86, replicates were pooled for subsequent analysis.

### Modelling global local RNA structures

Global RNA structure models for each variant were derived from SHAPE reactivity profiles and reference sequences through Superfold (v1.0) with a 600 nt maximum base pairing distance constraint and default SHAPE slope (1.8) and intercept (-0.6) parameters. ScanFold^41^ was used to estimate local fold stability with step size and window size parameters of 10 and 120 respectively, and 50 randomizations per step. Regions were ranked by Z-score where the lowest 20% corresponds to the highest (top 20%) stabilization values (Fig. 2a, Supplementary Fig. 4a). Generation of specific local and long-range structures was achieved with RNAstructure 6.4 modules partition-smp, Fold-smp, and stochastic-smp and the ViennaRNA 2.6.4 module RNAcofold.

### Identification of structurally conserved regions

Structure models were mapped to their respective aligned sequence to maintain consistency with multiple sequence alignment (MSA) numbering. Aligned structures were divided into 200 regions, each ≈150 bases long, to facilitate computing mean SHAPE-MaP reactivity, Shannon entropy, and a stability Z-score for each 150-nt block. Locations of the bottom 20^th^ percentile SHAPE reactivity, Shannon entropy, and stability blocks were identified in all 5 variants. Consensus regions for each metric were calculated as any region present in at least 3 out of 5 variants. Pearson correlation was also calculated in 150-nt windows for all 10 combinations of variant pairs. Any region measured to have an r value ≥0.8 was considered highly correlated. Consensus regions of highly correlated reactivity were designated as those present in 7 out of 10 structure pairs. Structure conservation was quantified as the fraction of identical base paired and unpaired positions within a specified 150-nt window between any two of ten variant pair combinations. Highly conserved structures are defined as those 20 regions demonstrating a contiguous structure conservation ratio of 0.85 or higher in at least 7 out of the 10 compared variants.

### Base pair covariance score calculation

The method for measuring base pair covariance was adapted from a procedure described by Sun *et al*.^12^ and updated to utilize the Infernal 1.1.5 software suite^42^ and a data set of 1,581 sequences of viruses belonging to *Coronaviridae* family which were culled for redundancy with CD-HIT^43^ from a larger database of 10,000 entries obtained at https://www.bv-brc.org. The cmbuild, cmcalibrate, and cmsearch modules were called from Infernal 1.1.5 in succession on each conserved structure in all variants, up to a maximum of 3 iterations or until no further sequences could be added to the covariance model. A covariance score for each structure within the 20 groups of conserved structures was calculated from the resulting alignment in accordance with published study^44^.

### Assessing the structural impact of SNVs

All SNV coordinates were gathered from the MSA and each was placed at the centre of a 100-nt window (±50 nt positions). Structure similarity at the corresponding genome coordinates of the two variants between which the SNV originates was then computed. The number of identical base (un)paired positions and mismatches were used to build a binary array of common (1) and distinct (0) identities. Arrays of a particular SNV type (A>N, U>N, G>N, C>N, N>A, N>U, N>G, N>C) were grouped into a 2D array and the mean value of each column was calculated as the overall structure conservation at that genome position.

### Measuring the effect of RBP interaction on viral RNA structure

Position weight matrices (PWMs) for human RBP motifs were taken from the ATtRACT database (ver. 0.99β). Each RBP PWM was used to scan all variant genome sequences for possible motif matches. Any matching motif coordinates were recorded as the genome positions of the respective variant. Differential gene expression (DGE) analysis was performed on variant pairs with DESeq2 (R Bioconductor), and gene names were obtained from Ensembl IDs using biomaRt. We cross-referenced the list of differentially expressed genes with RBP names from the PWM database and literature sources to identify which RBPs are significantly regulated. Analogous to the SNV analysis, the genome coordinates of significantly differentially expressed RBP motifs were analysed within a 100-nt window to compute aggregate structure conservation across all structure comparisons of a given RBP. RBPs with the largest potential impact on local genome structure were defined as those whose motifs experience low structure conservation (≤ 0.5) between a given VOC pair and significant changes in gene expression (|log2FC| ≥ 1 and padj < 0.05). Pairs of SARS-CoV-2 structure models of RBP motifs at regions of low structure conservation and significant DGE were modelled and reported.

### SPLASH analysis

Chimeric reads were divided into virus-virus and host-virus interactions for WT and Omicron XBB genomes. Virus-virus interactions were normalized to total virus-virus interactions and are shown in Fig. 4a (WT: blue, Omicron XBB: red). SPLASH hybrid structure models were generated with SHAPE-MaP reactivity as constrains using RNAcofold in the Vienna RNA package.

### LR1 secondary structure clustering

Scaling and clustering procedures were adapted from previous study^45^. One thousand LR1 structures were sampled from the Boltzmann ensemble with the stochastic-smp module from RNAstructure 6.4^46^ and folding free energy was calculated for all structures with efn2-smp. Duplicate structures were removed leaving N unique LR structures. Hamming base pair distances were calculated between all possible unique structure pairs to represent structural (dis)similarity as an N ⨯ N pairwise distance matrix. The scikit-learn multi-dimensional scaling (MDS) module was chosen to transform the distance matrix into two-dimensional embedding coordinates which are required for clustering analysis and plotting. The resulting embeddings were clustered with the HDBSCAN clustering algorithm. Optimal hyperparameter values were determined via grid search where the combination of values that maximized the Density Based Cluster Validity (DBCV) score were chosen for clustering.

### RNA A-to-I editing analysis

The A-to-I editing analysis pipeline was referred to previous study^27^ with some modifications. Raw reads were cleaned using FASTP^47^. Ten bases of 5’ and 3’ were trimmed and reads with more than 20% of unqualified bases were removed. The mean quality per base was fixed at a phred-score of 25. The reads that were shorter than 100-nt were discarded. The cleaned reads were mapped to the reference sequence using BWA^48^ aln (bwa aln-t 20) and mem (bwa mem-M-t 20-k 50). The unmapped reads were extracted for editing site calling as described in the published study^27^. Editing site calling was performed by subsampled each library to the same reads’ depth. To identify edited reads, read was specially required to have a minimum number of edited sites ≥ 4 (with either A-to-G or T-to-C sites in a read). The putative misaligned reads matching any of the criteria as shown in the study^27^ were also excluded. Only reads with the number of mismatch and low-quality A-to-G/T-to-C sites ≤ 2 were considered as edited reads. For each sample, the editing index was defined as number of editing sites from edited viral reads divided by the total number of viral reads in our study. The editing level was quantified by calculating the percentage of editing counts out of total reads at each position.

### Data analysis

Processing, analysis, and visualization of data were performed using python-3.11.7, R 4.3.3, and the associated modules numpy-1.26.4, scipy-1.12.0, scikit-learn-1.4.1, matplotlib-3.8.3, upsetplot-0.9.0, venn-0.1.3, seaborn 0.13.2, DESeq2-1.44.0, biomaRt-2.60.1, the stochastic-smp module from RNAstructure 6.4, and INFERNAL 1.1.5 modules cmbuild, cmcalibrate, and cmsearch. Calculation of statistical parameters of data sets (including means, medians, percentiles, and standard deviations) was performed using numpy. Statistical tests (Student’s t-tests) employed the appropriate functions in the scipy.stats module. Visualization was performed using matplotlib and seaborn for bar charts, pie charts, box plots, line plots, scatter plots, upset plots, Venn diagrams, and violin plots unless otherwise stated. *P*-values that were < 0.05 were considered statistically significant. *, *P* < 0.05, **, *P* < 0.01; ***, *P* < 0.001, ****, *P* < 0.0001.

## DATA AVAILABILITY

The data sets generated and analysed in this study are available in the GEO repositories, GSE279203 (SHAPE-map) and GSE285533 (SPLASH). The raw mass spectrometry spectra and search data were uploaded to the jPost repository^49^ with the following accession numbers: JPST003909 (jPOST), PXD065815 (ProteomeXchange). The authors declare that all other data supporting the findings of this study are available within the article and its Supplementary Information files or are available from the authors upon request.

## ACKNOWLEDGEMENT

We thank Dr. Meng How Tan for productive discussions, members of the Wan laboratory, A*STAR, for their input and suggestions during the course of this work. YW is supported by funding from A*STAR, National Research Foundation of Singapore, and NRF grant NRF-CRP27-2021-0003. AM is the recipient of program grant (PRG1154) from Estonian Research Council.

## AUTHOR CONTRIBUTIONS

YW, RH, AM, SLY came up with the project. YW, RGH, AM, SLY designed the experiments and the analysis together. SLY, SW performed most of the experiments. LD performed most of the analysis. LFW, JC, YHW, CKM and SW designed and performed the experiments that were carried out in BSL-3 laboratory. JSX performed ADAR1 eCLIP experiment. RS designed and performed the mass spectrometry experiments. KYT, SYL, ZZ supported the experiments. JH, YZ and JL supported the analysis. YW, RGH and SLY wrote the manuscript with everyone’s feedback.

